# 3,3′-Diindolylmethane Dose-Dependently Prevents Advanced Prostate Cancer

**DOI:** 10.1101/612929

**Authors:** Yufei Li, Nathaniel W. Mahloch, Nicholas J.E. Starkey, Mónica Peña-Luna, George E. Rottinghaus, Kevin L. Fritsche, Cynthia Besch-Williford, Dennis B. Lubahn

**Affiliations:** Department of Biochemistry, University of Missouri, Columbia, MO 65211, USA; MU Center for Botanical Interaction Studies, University of Missouri, Columbia, MO 65211, USA; Instituto Nacional de Medicina Genómica, Mexico City, 01900, Mexico; Veterinary Medical Diagnostic Laboratory, University of Missouri, Columbia, MO 65211, USA; Department of Nutrition and Exercise Physiology, University of Missouri, Columbia, MO 65211, USA; IDEXX BioResearch, 4011 Discovery Drive, Columbia, MO 65201

**Keywords:** 3,3′-Diindolylmethane, prostate cancer, TRAMP, weight loss, estrogen receptor

## Abstract

3,3′-Diindolylmethane (DIM) is an acid-derived dimer of indole-3-carbinol, found in many cruciferous vegetables, such as broccoli, and has been shown to inhibit prostate cancer (PCa) in several *in vitro* and *in vivo* models. We demonstrated that DIM stimulated both estrogen receptor alpha (ERα) and estrogen receptor beta (ERβ) transcriptional activities and propose that ERβ plays a role in mediating DIM’s inhibition on cancer cell growth. To further study the effects of DIM on inhibiting advanced PCa development, we tested DIM in TRAMP (TRansgenic Adenocarcinoma of the Mouse Prostate) mice. The control group of mice were fed a high fat diet. Three additional groups of mice were fed the same high fat diet supplemented with 0.04%, 0.2% and 1% DIM. Incidence of advanced PCa, poorly differentiated carcinoma (PDC), in the control group was 60%. 1% DIM dramatically reduced PDC incidence to 24% (p=0.0012), while 0.2% and 0.04% DIM reduced PDC incidence to 38% (p=0.047) and 45% (p=0.14) respectively. Though DIM did affect mice weights, statistical analysis showed a clear negative association between DIM concentration and PDC incidence with p=0.004, while the association between body weight and PDC incidence was not significant (p=0.953). In conclusion, our results show that dietary DIM can inhibit the most aggressive stage of prostate cancer at concentration lower than previously demonstrated, possibly working through an estrogen receptor mediated mechanism.

## Introduction

Prostate Cancer is the most prevalent cancer among U.S. males, other than skin cancer. It is estimated that in 2018 about 164,690 new prostate cancer cases will be diagnosed and that 29,430 patients will die from this disease, making it the second leading cause of cancer death in men [1]. Aging is the key factor for prostate cancer development and about one man in nine will get prostate cancer during his life time. Diet is another factor that can impact prostate cancer development. Epidemiological data has demonstrated that men in the U.S. and other developed countries, who usually have a western style high fat diet, have a much higher risk for prostate cancer [2]. Japanese and Chinese men have an increased risk for prostate cancer, after they emigrate and adopt a more western diet [3,4]. Though there is high incidence for prostate cancer in the U.S., the majority (91%) of them will be diagnosed as a localized cancer. The 5-year relative survival rate in a patient with local cancer is nearly 100%. However, the 5-year survival rate for metastatic prostate cancer is only 30% [5]. Our goal in this study was to assess the ability of DIM to prevent advanced prostate cancer.

Epidemiological studies have demonstrated that intake of cruciferous vegetables, such as broccoli, kale and cabbage will reduce the risk of advanced prostate cancer [6–8]. Indole-3-carbinol (I3C) is present at relative high levels in cruciferous vegetables and has been shown to have anti-cancer effects [9,10]. However, I3C is not stable at low pH and will spontaneously form 3,3′-Diindolylmethane (DIM) in the stomach. DIM is an acid-derived dimer and is considered to be the bioactive product of I3C [11].

DIM has been widely used as a dietary supplement in recent years and has been shown to have great therapeutic potentials for many diseases. There are more than ten clinical trials of DIM on different types of diseases, including prostate cancer, breast cancer, thyroid disease, cervical dysplasia, and others [12–16]. Research on this plant-derived compound have found that DIM interacts with several pathways, such as the estrogen receptor signaling pathway, aryl hydrocarbon receptor pathway, androgen receptor-signaling pathway, Nrf2 pathway and NF-κB pathway, suggesting DIM is a multi-target drug [17–23].

DIM has been shown to inhibit prostate cancer in several *in vitro* and *in vivo* models [21,24–29]. Wu. T. Y et al. previously showed that 1% dietary DIM prevented advanced prostate cancer (PDC) in the TRAMP model [21]. However, considering the hormesis effect, which shows an inverted U-shaped dose response curve, with only one concentration in the previous study, we could not decide which part of the response curve this concentration (1% DIM) represented [30–32]. This makes it difficult to provide concentration information for further preclinical and clinical studies. Additionally, DIM was reported to induce weight loss in mice fed with a high fat diet, which suggested DIM might be exerting its effect simply through weight loss [33]. In this paper, we tested DIM in the TRAMP model with 3 different concentrations: 0.04%, 0.2% and 1% w/w in a high fat diet to evaluate DIM’s effect on weight in parallel with advanced prostate cancer (PDC) prevention. Previously our lab demonstrated that in TRAMP mice, ERα stimulated advanced prostate cancer while ERβ inhibited it [34]. In this case, we examined DIM’s effect on estrogen receptor which suggested a role for ER in DIM’s prevention of advanced prostate cancer.

## Materials and Methods

### Reagents

17β-estradiol (E2) was purchased from Sigma Aldrich (St. Louis, MO). DIM used in cell culture was purchased from Sigma Aldrich (St. Louis, MO); DIM used in animal studies was purchased from BulkSupplements.com (Henderson, NV) and its purity was verified by HPLC as >98%. ICI 182,780 was a gift from Dr. Rex Hess (UI-UC).

### Animals

Animal study protocols were approved by the Animal Care and Use Committee of the University of Missouri and all institutional guidelines for animal welfare and use were followed. Study mice were bred by crossing male FVB TRAMP mice with C57BL/6 female mice to generate F1 C57/FVB hybrid male TRAMP mice as described previously [35]. TRAMP positive F1 males received study diets starting from 6 weeks of age, as TRAMP mice begin forming tumors after the onset of puberty (usually around 8 weeks of age) and continued until study termination. Study mice were housed 2-3 per micro-isolator cages and given free access to food and water, under the condition of a daily light: dark cycle of 12:12h, ambient temperature at 21°C, and humidity at 50% throughout the study. Mice were weighed every week to monitor body weight and tumor burden. At the age of 20 weeks, mice were euthanized, and their tissues were weighed and collected for further analyses. Prostates were fixed in 10% neutral buffered formalin and then embedded with paraffin. Tissue sections were stained with hematoxylin and eosin and were scanned by veterinary pathologists to determine cancer stages in a blinded fashion as we have previously described [34,35]. Prostates were staged as one of the following designations: 1) normal, 2) hyperplasia, 3) prostatic intraepithelial neoplasia (PIN), 4) well differentiated carcinoma (WDC), 5) moderately differentiated carcinoma (MDC) and 6) poorly differentiated carcinoma (PDC). Based on the findings by Bailman, et. al., PDC can arise independently from hyperplastic epithelial lesions and represents the primary malignant cells in the TRAMP model [36]. Because of these findings, if a small portion of the prostate had a poorly differentiated morphology, it was scored as PDC.

### Diet

High fat diet was purchased as pellets from Research Diets, Inc (New Brunswick, NJ), based on the Western Diet (D12079B) minus the usual added cholesterol (supplemental table 1). The fat content in the diet was 41 kcal%. There was a total of four groups of mice on study with 39-42 mice per treatment group. Except for the control, three doses of DIM, 0.04%, 0.2% and 1% w/w, were mixed and pelleted into the high fat diet by Research Diets, Inc (New Brunswick, NJ) as well. The highest 1% dose was chosen based on studies reported previously [21,37]. All diets were stored at −20°C to prevent oxidation and spoilage.

### Cell Culture

TRAMP-C2, DU145, LNCaP and MCF7 cells were obtained from the American Type Culture Collection (ATCC). U2OS cells were provided by Dr. Hansjorg Rindt (University of Missouri, Columbia, MO). TRAMP-C2 cells were maintained in Dulbecco’s modified Eagle’s medium (DMEM) (Invitrogen, Grand Island, NY) supplemented with 5% fetal bovine serum (FBS) (GE Healthcare Life Sciences, Logan, Utah). DU145, LNCaP cells were cultured in RPMI 1640 media (Invitrogen, Grand Island, NY) with 10% FBS. U2OS cells were cultured in phenol red-free DMEM/F-12 media (Invitrogen, Grand Island, NY) containing 5% FBS. MCF7 cells were maintained in DMEM (Invitrogen, Grand Island, NY) containing 10% FBS.

### Growth Assay

TRAMPC2, DU145, LNCaP and MCF7 cells were pre-incubated in 24-well/96-well plates in a phenol-red free, charcoal-stripped serum media for one day and then followed by treatment of 0-60uM DIM (0-90uM DIM were used for TRAMP-C2 cells). After treatment, cells were lysed with 1N NaOH after 0hr, 24hr, 48hr, 72hr and 96hr of additional growth. Total protein concentration was measured with the Bio-Rad DC kit (Bio-Rad, Hercules, CA) by measuring absorbance at 750nm. Total protein had been previously shown to correlate with the thymidine incorporation assay in prostate cancer cells, as a reliable measure for cell growth [38].

### Luciferase Assay

U2OS cells were pre-incubated in phenol red-free DMEM/F-12 media supplemented with 5% charcoal-stripped FBS. After 24 hours, cells were transfected with the vitellogenin ERE luciferase reporter plasmid, the SV40-Renilla control vector, and either the ERα or ERβ expression vector using Fugene HD (Promega, Madison, WI) for 18 hours. Transfected cells were seeded to 96-well plates and then followed by treatment of 0-30uM DIM and DIM with 100nM ICI 182,780 for 24 hours. After that, cells were lysed using passive lysis buffer (Promega, Madison, WI) and transferred to assay plates. Luciferase and Renilla activities were measured by Biotek Synergy 2 plate reader (Winooski, VT) using the Dual-luciferase assay kit (Promega, Madison, WI).

## Results

### DIM stimulates both estrogen receptor alpha (ERα)and estrogen receptor beta (ERβ) transcriptional activities

We examined the effects of DIM on ER’s transcriptional activity by using the *in vitro* luciferase transcription assay. Human Bone Osteosarcoma Epithelial Cells (U2OS) were cotransfected with the vitellogenin estrogen response element (vERE) driven luciferase reporter, the SV40-Renilla control vector, and expression vectors for ERα or ERβ. The transcriptional activity was determined by measuring luciferase expression over renilla expression. DIM activated both ERα and ERβ transcriptional activities in a dose dependent manner. EC50 of ERα was ∼3uM (Figure 1A), while EC50 of ERβ was ∼10uM (Figure 1B). DIM’s stimulation of the transcriptional activities of both receptors could be abolished by adding the ER antagonist ICI 182,780, indicating these activations are mediated by estrogen receptors. Similar effects were also observed in MCF7-derived ER negative C4-12-7 cell lines (Supplemental Figure 1), indicating these are not cell-type specific effects.

**Figure 1.**
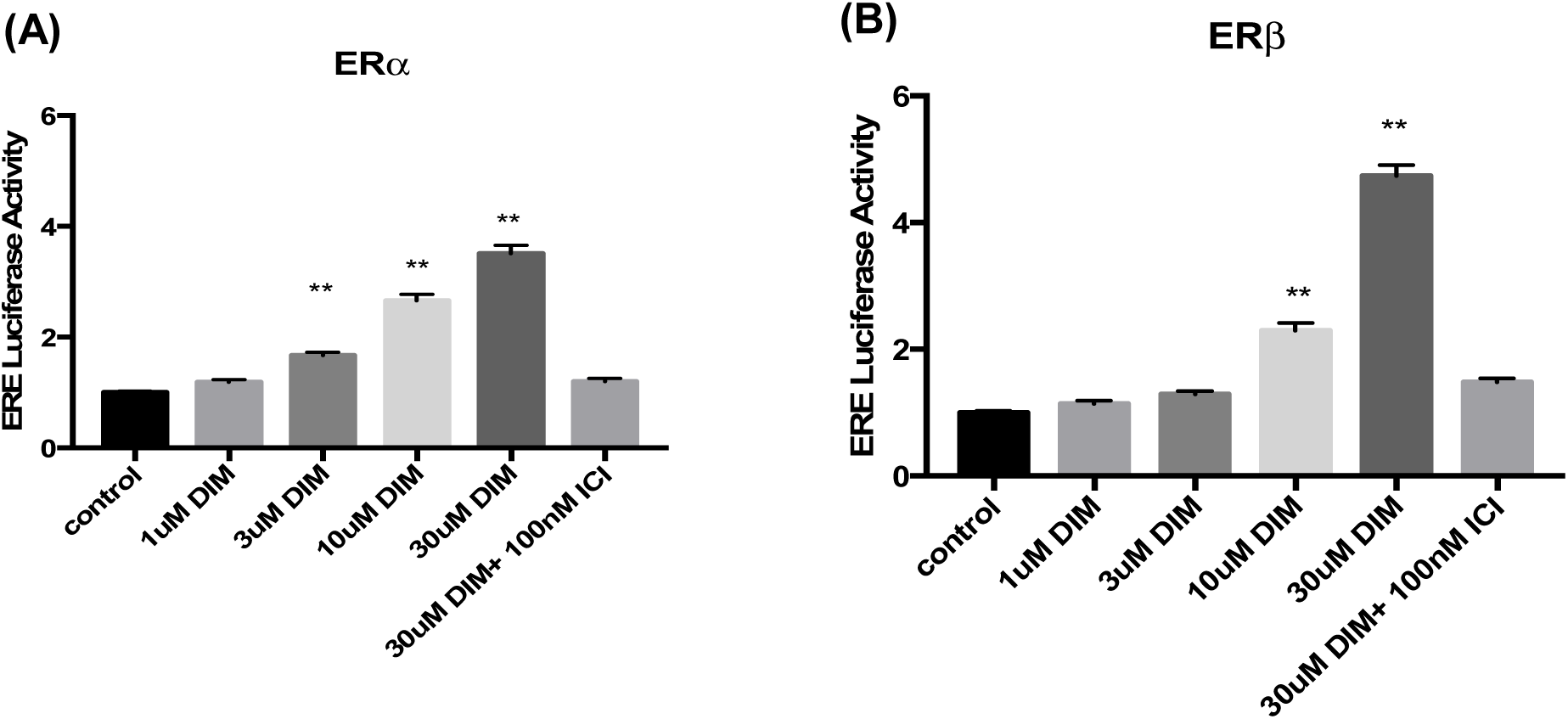
DIM stimulates ERa **A)** and ERβ **B)** transcriptional activities in an ERE luciferase assay in U2OS cell. Error bars indicates S.E.M. ** indicates p<0.001 compared to control

### DIM inhibited the growth of ERβ positive cancer cell lines

The two nuclear estrogen receptors, ERα and ERβ, play opposing roles in cell growth. ERα is proposed to be involved in proliferative process, while ERβ usually shows anti-proliferative effects and increases apoptosis [34,39]. We tested DIM for its effects on cell growth in five cell lines each with a different ER status. Prostate cancer cell lines LNCaP and DU145 only express ERβ and not ERα [40]. Cells were treated with or without DIM at three concentrations: 10uM, 30uM and 60uM for 96h. DIM inhibited the growth of LNCaP and DU145 at all three concentrations (Figure 2A, B). Prostate cancer cell line PC3 expresses both ERα and ERβ [40]. The inhibitory effects of DIM on cell growth were only observed at higher concentrations, while low concentration (10uM) of DIM showed no effects (Figure 2C). In contrast, in MCF7 cell line, a human breast cancer cell line which only expresses ERα, 10uM DIM stimulates the cell growth (Figure 2D). This is consistent with the observation from another group that low concentrations of DIM stimulated ERα and induced cell proliferation [41]. In addition, the MCF7 cells were more sensitive to DIM, with 60uM DIM showing cell toxicity. TRAMP-C2 is a mouse prostate cancer cell line derived from TRAMP mice that expresses both ERα and ERβ (supplemental figure 6). DIM inhibited the growth of TRAMP-C2 with IC50 >30uM (Figure 2E). Taken together, our results suggested that ERβ might play an important role in mediating the inhibitory effect of DIM on cell growth.

**Figure 2.**
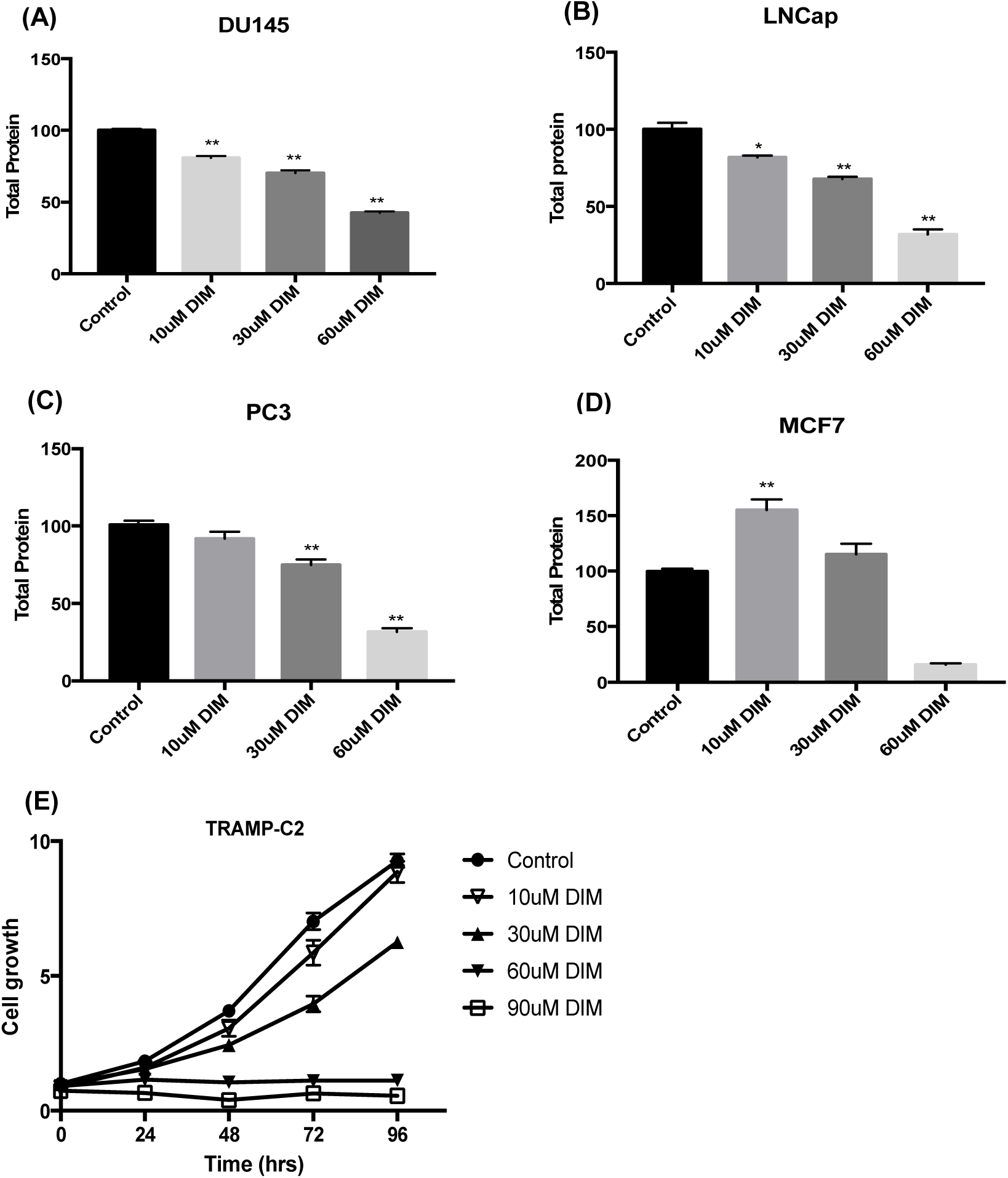
Effects of DIM on cell growth. **A)** Du145, **B)** LNCaP, **C)** PC3 and **D)** MCF7 cells were plated in 24-well plate at 50% confluence. Cells were then treated with 0-60uM DIM for 96 hrs. The total cellular protein concentration was measured and normalized to the control of each cell line. **E)** TRAMP-C2 cells were plated in 24-well plate and treated with 0-90uM DIM. The total cellular protein concentrations were determined every 24hrs for total 96hrs. Protein concentrations were normalized to the control at 0h. Error bars indicates S.E.M. * indicates p< 0.01, ** indicates p<0.001 compared to control

### DIM prevented advanced prostate cancer development in TRAMP mouse

Study mice were placed on a high fat diet to mimic the western style high fat diet among U.S males. The diet of the three experiment groups were supplemented with 1%, 0.2% and 0.04% (w/w) DIM. Incidence of PDC, which is the most aggressive stage of prostate cancer in TRAMP mouse, in the control group was 60%. By adding DIM to the diet, we saw a clear trend line of decreasing PDC incidence, with 0.04%, 0.2% and 1% DIM reducing PDC incidence to 45% (P=0.14), 38% (P=0.047) and 24% (p=0.0012), respectively (Table 1). While an increase in Well Differentiated Carcinoma (WDC) occurred in mice treated with 1% DIM (p= 0.028), the overall prevalence of cancer, which included WDC, MDC and PDC, did not change between groups. 1% DIM had a slightly lower cancer incidence of 91% and it was not significant from other groups. (Table 1).

**Table 1.** Effects of DIM on the incidence of prostate cancer in 20-week-old TRAMP mice. DIM inhibited PDC incidence in a dose-dependent manner. The inhibitory effect of 0.04% DIM was not significant (p=0.14). Both 0.2% DIM (p= 0.047)^a^ and 1% DIM (p=0.0012)^b^ had a statistically significant effects on PDC when compared to the control. There was also an apparent trend line of increasing WDC incidence, but only 1% DIM had a statistical effect (p=0.028)^c^. Incidence of tumor stage were analyzed using Fisher’s exact one-tailed *P* value.

| Group | N | Tumor Stage |  |  |  |  |  |
| --- | --- | --- | --- | --- | --- | --- | --- |
|  |  | Non-cancer<br>n (%) |  |  | Cancer<br>n (%) |  |  |
|  |  | Normal | Hyperplasia | PIN | WDC | MDC | PDC |
| Control | 42 | 0 | 0 | 0 | 17 (40%) | 0 | 25 (60%) |
| 0.04% DIM | 42 | 0 | 0 | 0 | 23 (55%) | 0 | 19 (45%) |
| 0.2% DIM | 39 | 1 (3%) | 0 | 0 | 23 (59%) | 0 | 15 (38%) <sup>a</sup> |
| 1% DIM | 41 | 0 | 1 (2%) | 3 (7%) | 27 (66%) <sup>c</sup> | 0 | 10 (24%) <sup>b</sup> |

In our study, we did observe an effect of DIM on mouse body weight in mice fed with 1% DIM (Figure 3). At age 20 weeks, mice in the 1% DIM group had a lower average body weight of 29.5g compared to the control (34.9g) (Figure 3B). During the weekly monitoring of weight, we found that mice treated with 1% DIM had a dramatic decrease in body weight, after the first week of treatment. Though they seemed to recover at the second week, they were never able to catch up with the other groups (Figure 3C). Epidemiological studies have suggested that obesity positively associates with the risk of aggressive prostate cancer [42,43]. In order to eliminate any possible confounding effect of body weight on PDC incidence, we fit our data to a logistic regression using both body weight and dosage of DIM as the independent variables and PDC incidence as the dependent variable. The result of the logistic regression indicated that the incidence of PDC decreased as we increased the dose of DIM, even controlling for body weight. The association between PDC incidence and DIM dosage was statistically significant (p=0.004), while the association between PDC incidence and body weight was not significant (p=0.953) (Figure 4).

**Figure 3.**
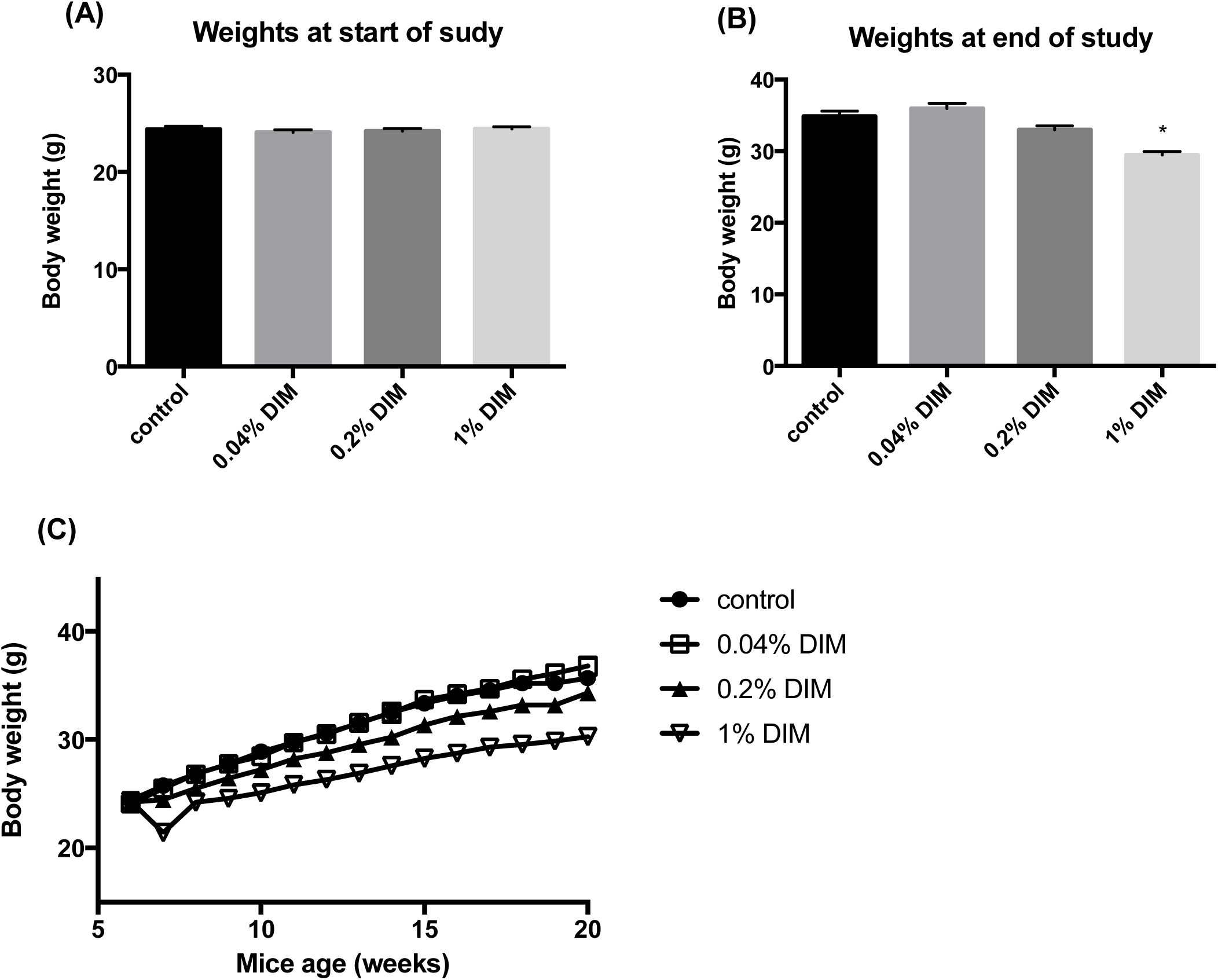
Effect of DIM on average mouse body weight. **A)** Average body weight of each group at the beginning of the study. **B)** Average body weight of each group at the end of the study. **C)** Time course change of the body weight. All mice on study were measured weekly and average body weight was calculated for each group. Error bars indicates S.E.M. * indicates p< 0.01

**Figure 4.**
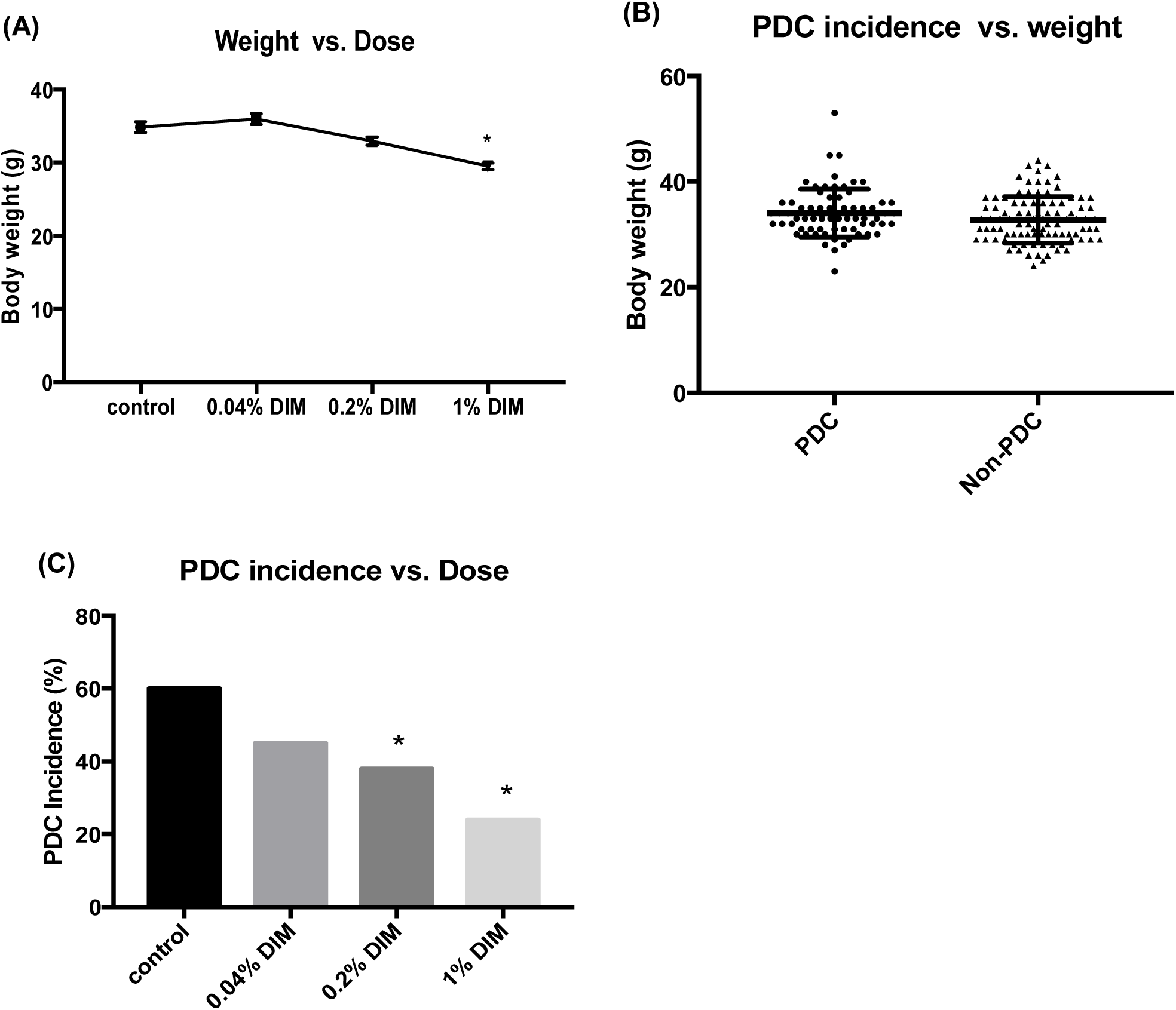
Correlation between mouse body weight, dosage of DIM in the diet and PDC incidence. Mice body weights were measured at the end of study (age of 20 weeks). **A)** Average mouse body weight decreased as increasing DIM concentration in the diet. Error bars indicates S.E.M. **B)** Average body weights of mice with PDC vs. without PDC. Error bars indicates S.D. **C)** Increasing DIM concentration in the diet decreased PDC incidence. Error bars indicates S.E.M. * indicates p< 0.01

## Discussion

DIM has been reported by Vivar et.al to be a selective ERβ agonist in U2OS, HeLa and MDA-MB-231 cell lines in the luciferase reporter assay and does not stimulate ERα at all [17]. Interestingly, we did find DIM activated both estrogen receptors in U2OS and C4-12-5 breast cancer cell lines. Our data is in agreement with several other reports that DIM activates ERα in MCF7 breast cancer and Ishikawa cell lines either using luciferase reporter assay or measuring the endogenous ERα- regulated gene expression [41,44].

On the other hand, DIM does not compete with E2 in a radio-labeled E2 competitive binding assay (Supplemental figure 2) [17,45]. The explanation of this ligand-independent activation of DIM is still not clear. One explanation is that DIM activates ERβ by recruiting ERβ coactivators, since silencing of SRC-2 can inhibit the activation of ERβ targeted keratin 19, NKG2E, and CECR6 genes by DIM [17]. Other groups also reported protein kinase A (PKA) inhibitor H89 ablated DIM’s activation of ER, indicating PKA ‘s involvement and phosphorylation of ER during this activation [41,44,45].

Additionally, DIM is reported to be a weak agonist of Aryl Hydrocarbon Receptor (AhR) [46,47]. When AhR is activated, it translocates into the nucleus where it forms a complex with the AhR nuclear translocator (Arnt). This heterodimer complex can directly associate with ERα and ERβ, recruiting the co-activator p300, which then results in ligand independent activation of estrogen receptors [48]. Recently, our lab reported an alternate binding site present in ERβ, which can allosterically regulate ERβ’s activity [49]. It is possible that DIM binds and activates ER from an alternate binding site but does not allosterically influence the binding affinity of E2 in the orthosteric site in an appreciable way. Thus, DIM is not a competitive ligand with estradiol on estrogen receptors.

From our published data, we have shown that knocking out ERβ in transgenic mice results in almost double the incidence of PDC in the TRAMP mouse model, while knocking out ERα results in almost no advanced prostate cancer [34]. Because of the opposing biological functions of ERα and ERβ in cancer, we performed growth assays in cell lines with different ER subtypes to determine if estrogen receptor mediates the inhibitory effect of DIM on cancer cell lines. DIM only showed dose-dependent inhibitory effect in cell lines with ERβ present (Figure 2 A-C, E). In ERα-only MCF7 cells, DIM even showed a stimulation of growth at 10uM (Figure 2D). Interestingly, at a higher 30uM DIM concentration, the growth stimulation was decreased. This suggests more than one pathway is regulated by DIM in the MCF-7 cell line.

We observed an inhibitory effect of DIM on the growth of TRAMPC2 cells in the concentration range between 10 to 90uM. It is reported that following a one-time gavage consumption of 250mg DIM/kg in mice, the physiological concentration of DIM in plasma ranged from a high of 16uM to a low of ∼ 0.3uM over 24 hours [37]. In our study, the mice in the 1% DIM diet ate about 4 times more DIM per 24 hours than in the reported gavage study. Therefore, our *in vivo* and *in vitro* studies were performed in overlapping concentration ranges.

There are a few reports of DIM inhibiting prostate cancer in TRAMP mice [21,27–29]. Wu. T. Y et al. reported that 1% DIM in diet inhibited advanced prostate cancer [21]. However, the inverted U shape of response vs. concentration phenomenon is common with hormone receptors responses [30–32]. In order to test the effect of DIM in the concentration range we used *in vitro,* and also to check if it would follow the same dose dependent inhibition pattern as we observed *in vitro,* we tested DIM *in vivo* at three different doses: 0.04%, 0.2% and 1% in diet. DIM very nicely followed the pattern, dose-dependently inhibiting PDC incidence (Table 1).

The increase in WDC incidence was surprising initially. However, this increase is not real, but instead is just the bias of our method for scoring the cancer stage. PDC arises independently of epithelial changes, and in our experience, small foci of PDC expand to involve all lobes of the prostate given enough time. Using the standard staging methodology, identification of PDC in any prostate lobe, regardless of the status of epithelial hyperplasia or neoplasia in the dorsolateral prostate lobe, is scored as PDC.

Thus, a prostate with a mixture of a large portion of WDC but a small focus of PDC is determined to be PDC. Based on this approach, it is possible that the overall incidence of WDC is not changed, but rather that PDC incidence is decreasing. There is no statistical difference in the overall incidence of cancer (WDC, MDC and PDC combined together) between all treatment groups. Thus DIM decreased the incidence of PDC, which might reveal the WDC that used to be “covered” by PDC, and there just seemed to be an increase in WDC incidence. A possible future solution to decrease WDC would be to combine DIM and another compound that inhibits WDC in the diet, for example, genistein, which we showed previously only inhibited WDC and had no effect on PDC [34].

Several previous studies of DIM in TRAMP mice did not report the influence of DIM on body weight [21,27–29]. There are two reasons that may explain the difference between our results and these previous reports. First, they used low fat diets which contained less than 10% w/w fat in the diet. In contrast, we used a high fat diet that consisted of 21% w/w fat. DIM is reported to suppress obesity when fed with a high fat diet [33]. Second, the genetic background of our study mice was different than others. We used the C57/FVB F1 hybrid TRAMP mouse, while they used C57BL/6 TRAMP mice.

We demonstrated that DIM stimulated estrogen receptors. Because ERβ is reported to alleviate obesity [50], we wanted to know whether the initially effect of DIM on body weight was mediated by ERβ or simply by not eating because mice did not like the taste of DIM. Two additional experiments were performed: 1) Feeding mice with different ER genotypes (both ERα and ERβ wild type, hetero and knockout) on control or 1% DIM diet, then comparing the body weight change (supplemental Figure 3); 2) Treating mice with different ER genotypes by injecting either corn oil or 200mg/kg DIM (supplemental Figure 4). It turned out that the body weight change was observed in all mice that we fed with 1% DIM but not in controls, regardless of the ER genotypes. On the other hand, injection of DIM in corn oil had no influence on the body weight. Additionally, the average daily food intake of mice in the 1% DIM group was much lower than that of control, when measured during the first two weeks on study (supplemental figure 5). Altogether, this suggests that the initial drop of body weight of mice in the 1% DIM group is not related to the estrogen receptor, but likely because of the taste of the diet which lowers the food intake.

In conclusion, we demonstrated that DIM stimulated both estrogen receptors, and that ERβ played an important role in mediating the inhibition of DIM on cancer cell growth. DIM dose-dependently prevented advanced prostate cancer development when fed high fat diets. Future studies to confirm ER’s importance in DIM’s function, including double transgenic ER knockout/TRAMP mice studies will need to be performed.

## Acknowledgment

We would like to thank Dr. Hansjorg Rindt for providing the U2OS cell line. We also appreciate the help of Peggy A. Eichen in the TRAMP study. We would like to thank the Mexican National Council for Science and Technology (CONACYT) for the fellowship (CVU 630757) to support the work of Mónica Peña-Luna on this project. This work was supported by grants from the Department of Defense (DOD) – Congressionally Directed Medical Research Programs (CDMRP) - PC100455, and the National Center for Complementary and Integrative Health (NCCIH), the Office of Dietary Supplements (ODS), and the National Cancer Institute (NCI) - P50AT006273. The contents herein are solely the responsibility of the authors and do not necessarily represent the official views of the DOD, NCCIH, ODS, NCI, or the National Institutes of Health.

