## Supplemental data for "3,3′-Diindolylmethane Dose-Dependently Prevents Advanced Prostate Cancer"

**Advanced Prostate Cancer**

**Supplemental table 1**: Formulation of high fat diet

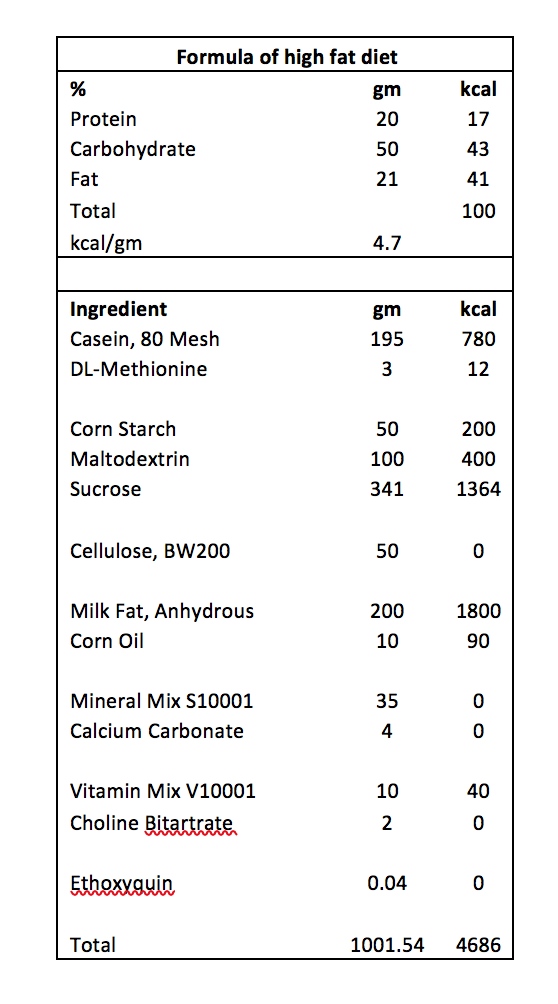

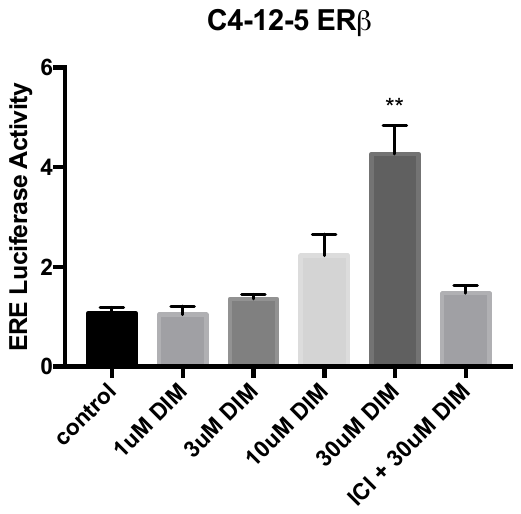

**(B)**

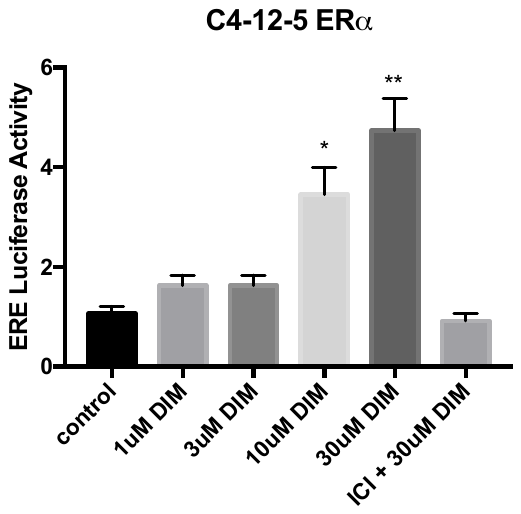

**(A)**

**Supplemental figure 1.** DIM stimulates ERa **A)** and ERβ **B)** transcriptional activities in an ERE luciferase assay in MCF7 derived ER negative C4-12-7 cells. Error bars indicates S.E.M. * indicates p< 0.01, ** indicates p<0.001 compared to control

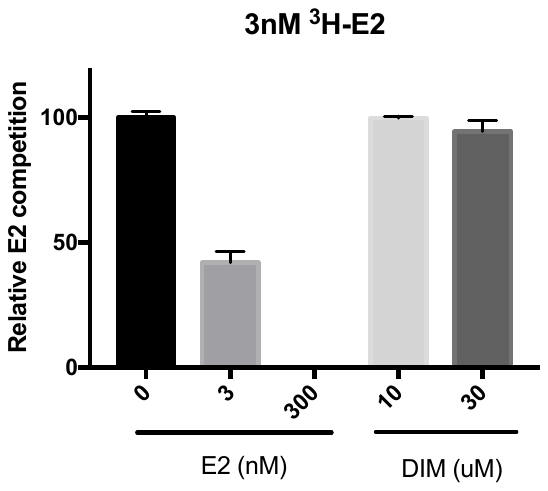

**Supplemental figure 2.** Competitive binding of DIM with human estrogen receptor hERβ. DIM does not compete with tritiated estradiol (^3^H-E2) (3nM) for binding to hERβ but in the positive control the unlabeled E2 does. Error bars indicates S.E.M.

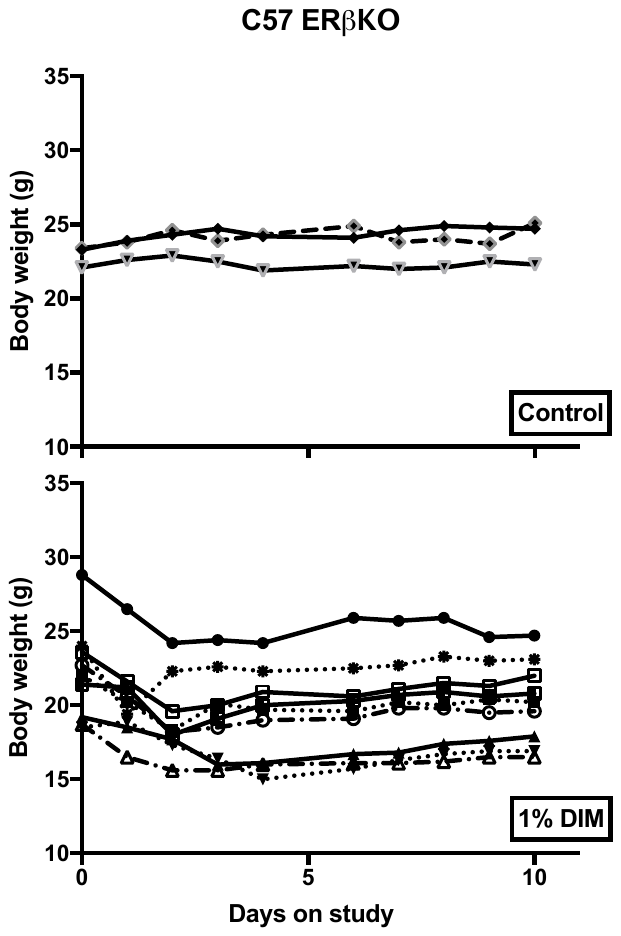

**(A)**

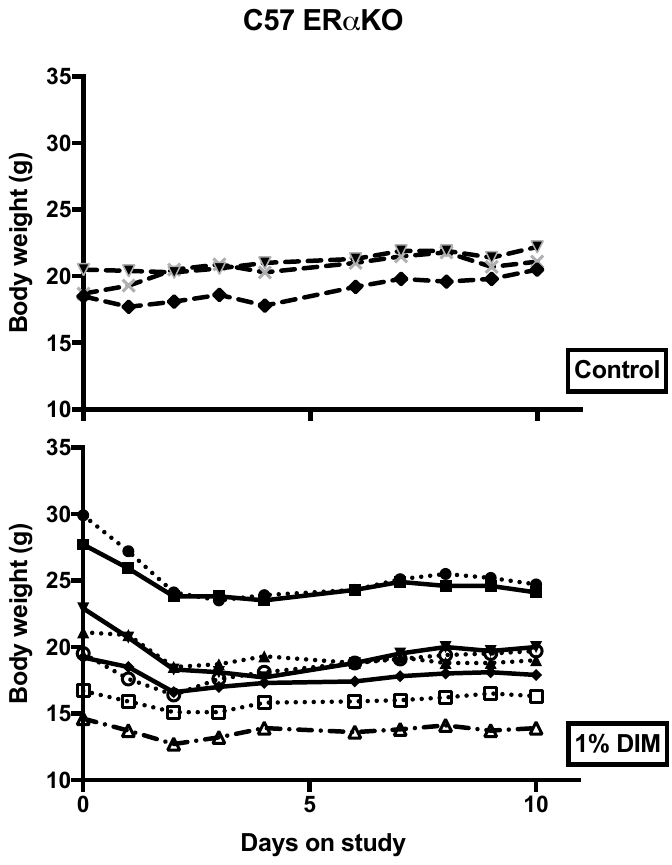

**(B)**

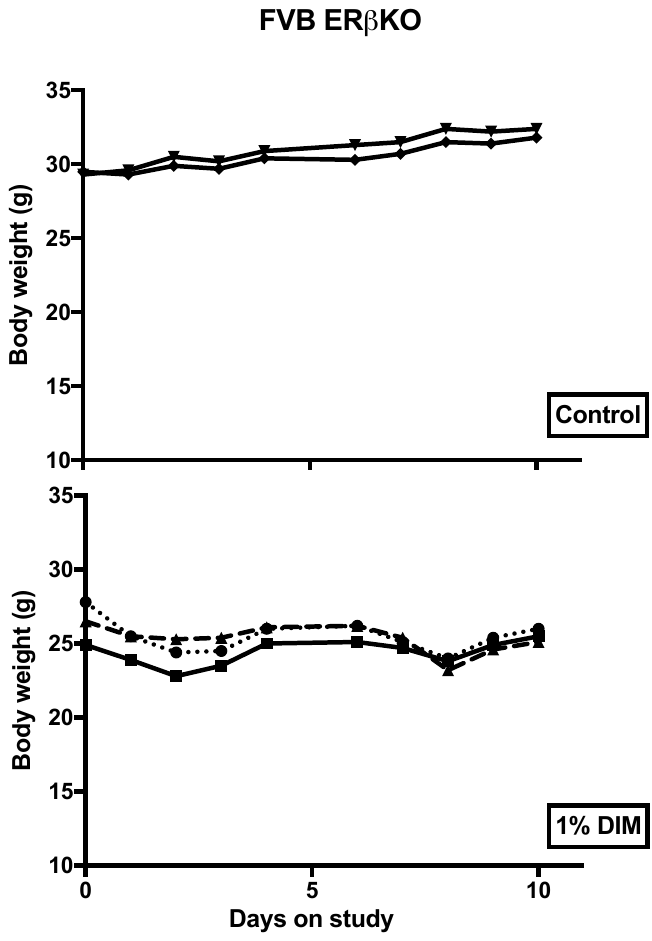

**(C)**

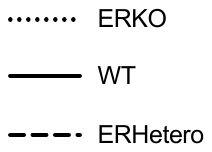

**Supplemental figure 3.** Time course body weight change of mice on control or 1% DIM diet. Mouse body weight was measured every day for 10 days. Each line represents one mouse. Solid line (**－**) is wild type mouse; dashed line (--) is ER hetero genotype mouse and dotted line (**^......^**) is ER knockout mouse.

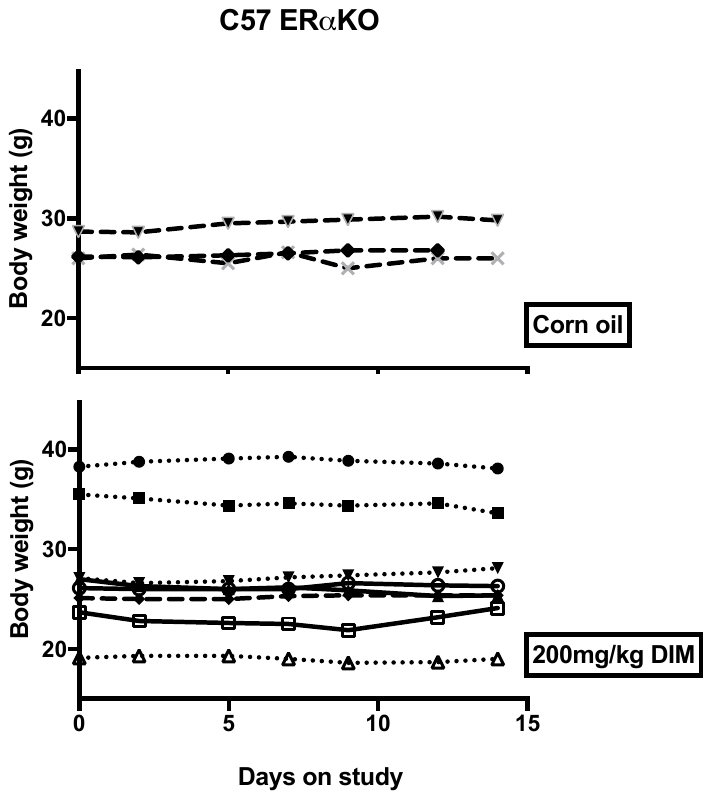

**(B)**

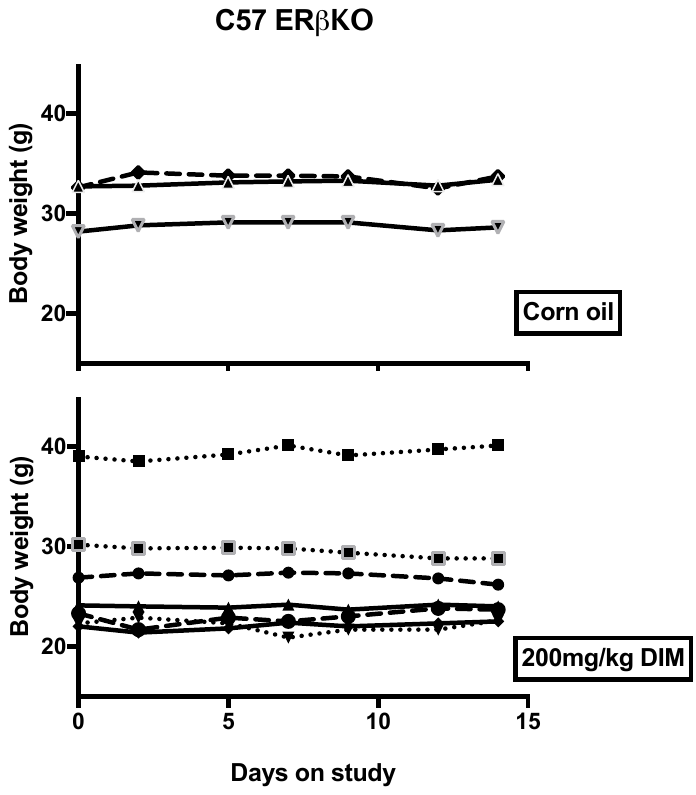

**(A)**

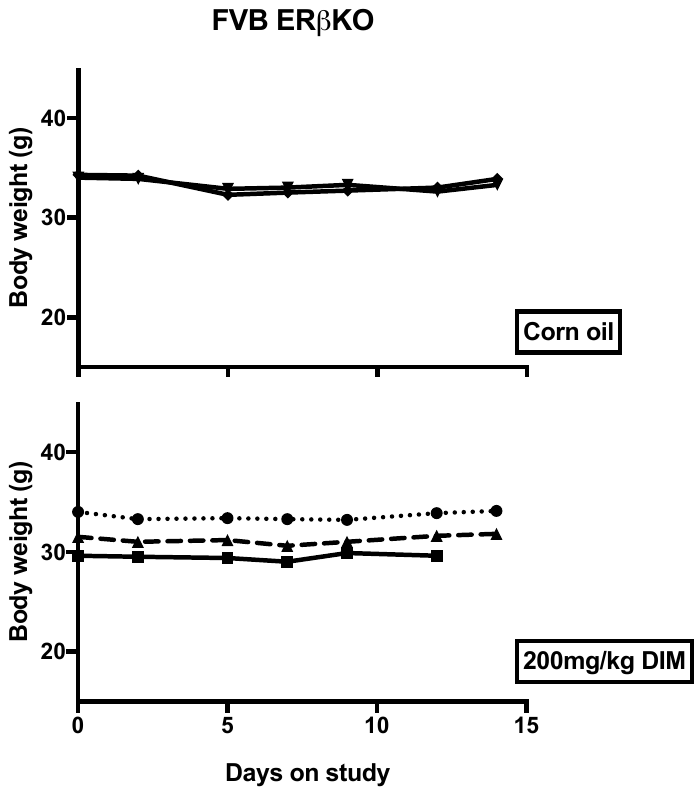

**(C)**

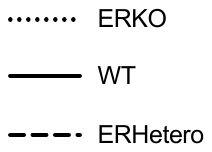

**Supplemental figure 4.** Time course body weight change of mice intraperitoneal

injected with corn oil or 200mg/kg DIM. Mouse body weight was measured every day for 14days. Each line represents one mouse. Solid line (**－**) is wild type mouse; dashed line (--) is ER hetero genotype mouse and dotted line (**^......^**) is ER knockout mouse

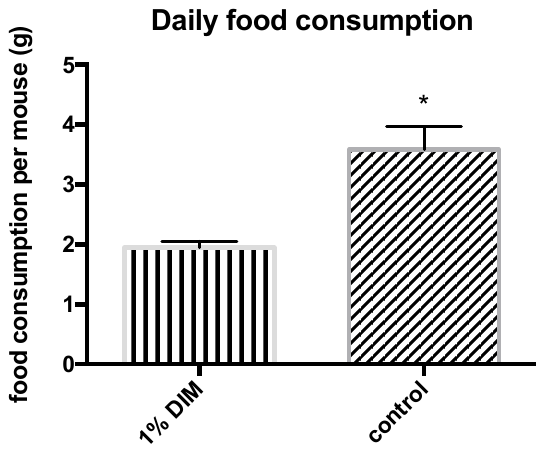

**Supplemental figure 5.** Average daily food consumption of control (N=9) and 1% DIM (N=17) treated mice in the ERKO feeding study. Diets were provided as pellets and placed in a jar that allowed mice to climb into the jar to get the food. A wire entanglement was placed above the food to prevent mouse taking the food pellets out of the jar. The food jar was measured daily to calculate the average daily consumption

of food over a two-week period. Error bars indicates, S.E.M. * indicates p< 0.01.

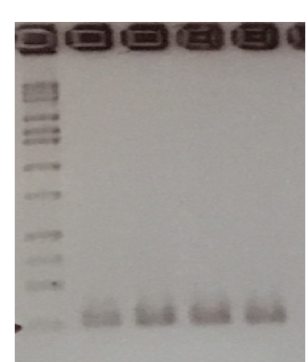

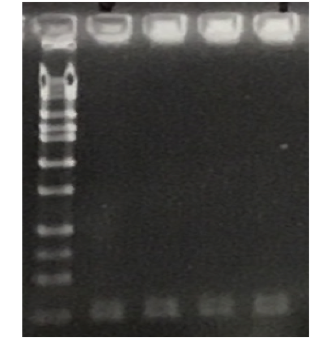

**ER𝚊**

**ERβ**

1	2

1	2

**Supplemental figure 6.** RT-PCR analysis of ERa and ERβ expression in TRAMP-C2 cell line. Lane 1 and 2 are duplicates of amplified cDNA libraries made from 2 independent mRNA samples. Primer sequences for ERa: fERa- GACCAGATGGTCAGTGCCTT (1139-1158); rERa- ACTCGAGAAGGTGGACCTGA. (1343-1324). Primer Sequences for ERβ: fERβ- GTAGAGAGCCGTCACGAATACT (166-187); rERβ- GGTTCTGCATAGAGAAGCGATG (362-341). Arrows indicate presence of 205bp ERa and 197bp ERβ bands.
